# An open-source tool to assess the carbon footprint of research

**DOI:** 10.1101/2021.01.14.426384

**Authors:** Jérôme Mariette, Odile Blanchard, Olivier Berné, Olivier Aumont, Julian Carrey, Anne Laure Ligozat, Emmanuel Lellouch, Philippe-e Roche, Gäel Guennebaud, Joel Thanwerdas, Philippe Bardou, Gérald Salin, Elise Maigne, Sophie Servan, Tamara Ben-Ari

## Abstract

The scrutiny over the carbon footprint of academics has increased rapidly in the last few years. This has resulted in a series of publications providing various estimates of the carbon footprint of one or several research activities, principally at the scale of a university or a research center or, more recently, a field of research. The variety of tools or methodologies - on which these estimates rely - unfortunately prevents from any direct comparison because of the sensitivity of carbon footprint assessments to variations in the scope and to key parameters such as emission factors. In an effort to enabling a robust comparison of research carbon footprints across institutions, contexts or disciplines, we present an open-source web application, *GES 1point5* designed to estimate the carbon footprint of a department, research lab or team in any country of the world with a transparent and common methodology. The current version of *GES 1point5*, open-source and freely available, takes into account the most common and often predominant emission sources in research labs: buildings, digital devices, commuting, and professional travel. *GES 1point5* is developed by an interdisciplinary team of scientists from several public research institutions in France as part of the Labos 1point5 project. *GES 1point5* is therefore presently tailored for the French context but can be adjusted to any national contexts by adjusting the values of emission factors. The versatility and usability of the software have been empirically validated by its adoption by several hundred research labs in France over the last 18 months. In addition to enabling the estimation and monitoring of greenhouse gas (GHG) emissions at the scale of a research lab, *GES 1point5* is designed to aggregate the data entered by the labs and the corresponding GHG emissions estimates into a comprehensive database. *GES 1point5* can therefore allow to (i) identify robust determinants of the carbon footprint of research activities across a network of research labs (ii) estimate the carbon footprint of research at the national scale. A preliminary analysis of the carbon footprint of more than one hundred laboratories is presented to illustrate the potential of the approach. While assessments of carbon footprints are often externalized onto extension services and proprietary softwares, *GES 1point5* is designed as a hands-on, pedagogic and transparent tool for research labs to monitor and reduce their own carbon footprint. This internalization has strong positive co-benefits for academics in terms of awareness and empowerment. We further expect that international dissemination of *GES 1point5* will contribute to establishing a global understanding of the drivers of the research carbon footprint worldwide and an identification of the levers to decrease it.

**Availability and implementation:** *GES 1point5* is available online at http://labos1point5.org/ges-1point5 and its source code can be downloaded from the GitLab platform at https://framagit.org/labos1point5/l1p5-vuejs.

## 1 Introduction

Drastic reductions of GHG emissions are needed to bend current global emission rates. After signing the Paris Agreement, many countries including France, have committed to reach carbon neutrality by 2050. These commitments imply the implementation of very ambitious GHG mitigation strategies in all economic sectors. Although it may be expected that the direct contribution of the teaching and research sectors to national GHG emissions is relatively small, especially in comparison to other sectors of activities, academia is bound to contribute to these GHG reduction efforts for several important reasons. First, because of the role that academia plays in producing and imparting knowledge on climate change and its impacts on ecosystems and societies. Second, scientists contribute more and more actively to the public debate around climate change mitigation strategies. As a consequence, they are expected to act as role models for the sake of their “credibility” [Attari et al., 2016]. Third and finally, available assessments suggest that the carbon footprint of scientists is above the per-capita median value in their countries of residence, [Nature Astronomy, 2020, Spinellis and Louridas, 2013, Fox et al., 2009, Grémillet, 2008]. The work presented in this paper builds on these imperatives and thrives to contribute to making research a leading sector in the transition towards a low-carbon society.

At the international level, only a small number of studies assess the carbon footprints of the research sector. These studies tend to focus on one single entity such as a research department or a university [Güereca et al., 2013, Wynes et al., 2019], a specific event such as a conference [Spinellis and Louridas, 2013, Desiere, 2016, Stroud and Feeley, 2015, Klöwer et al., 2020], a single source of GHG emissions such as air-travel [Ciers et al., 2019], a specific research project [Achten et al., 2013, Barret, 2020, Aujoux et al., 2021] or specific instruments in a field of research [Knödlseder et al., 2022]. Most often, these assessments are only available for one single year [Güereca et al., 2013]. Therefore these studies present methodologies that are not designed to estimate a comprehensive carbon footprint for a wide variety of research laboratories over multiple years. To the best of our knowledge, two softwares have been developed for assessments at the university level, but they do not specifically address research activities : SIMAP^®^^1^ and CO2UNV [Valls-Val and Bovea, 2022].

Beyond the specificities of the research sector, several standards are widely used for carbon footprint assessment. The GHG Protocol^2^ offers international standards to account for GHG emissions at corporate, city or country levels. Some methodologies are deployed on a nationwide scale. For instance, in France, *Bilan Carbone*®^3^ supported by Association Bilan Carbone, offers a generic methodology (developed as a set of spreadsheets) which may be used at various scales. Unfortunately, *Bilan Carbone* requires its users to pay for a licence and a training program. Other tools, freely available, are designed to estimate individual or households’ GHG emissions and are unfit in the research context, e.g., [myclimate] ^4,5,6,7,8^.

When different methodologies are used by several entities, comparing their carbon footprints is virtually impossible since discrepancies in the results cannot be robustly attributed to differences in emissions. Comparisons are possible only at the price of *a posteriori* numerous data adjustments or assumptions Helmers et al. [2021]. Instead, sharing the same methodology, perimeter and parameters allows to consistently compare the results of a very large number of entities, and store footprints for large scale analyses.

Here, we present *GES 1point5* ^9^, a free and open-source standardized application to assess the carbon footprint of research activities conducted at the scale of a research group called a “laboratory”^10^. It is built as a web application - which is more user-friendly, operating system agnostic and interoperable than spreadsheets. The hypotheses, values and methodology are presented transparently and included in the tool. *GES 1point5* is developed by an interdisciplinary team of engineers and researchers from various research fields in France who interact in the *groupement de recherche (GDR) Labos 1point5*^11^. *Labos 1point5* gathers hundreds of scientists and research staff working together to estimate and mitigate the impacts of research activities on the environment.

More specifically, *GES 1point5* allows research labs to (i) estimate the emissions attributed to the energy consumption and refrigerant gases of their buildings, those attributed to the purchase of their digital devices, commuting, professional travel as well as the associated uncertainties, (ii) easily highlight, via a graphic interface, the drivers of their main GHG emissions, and (iii) design emission reduction actions and evaluate their mitigation effects over time. Because they rely on a standardized protocol, *GES 1point5* -based carbon footprints can be compared between research labs, disciplines, or contexts. This will, in turn, widely increase our understanding of prevailing emission sources within research activities and their heterogeneity (e.g., disciplinary, sociological, or geographic). At the national scale, the data collected will allow to estimate the carbon footprint of the public research sector and thus support the exploration of evidence-based emission reduction strategies. Notwithstanding the specific institutional context of the public research sector in France, *GES 1point5* is a standardized tool that may be replicated in any foreign research center with minimal adjustments (see Section 3.2). A French and an English version of *GES 1point5* are built-in in the current version to ease the application deployment in any country.

In the next sections, we first present the main features of the French research system, and the goals of *GES 1point5* ; we then explain the methodology (Section 3) and the implementation of *GES 1point5* (Section 4) together with an illustration of its outputs, namely the GHG inventory and the carbon footprint of a fictitious research lab(Section 5). To illustrate the potential of the present tool, a preliminary analysis of the typology of GHG inventories across 109 French research laboratories is also presented (Section 6). Future developments, research perspectives and conclusion are finally addressed.

## 2 Context and goals

The French research system encompasses several types of institutions ^12^: national research institutes (such as CNRS, INRAe, CEA, IRD, INRIA …), semi-public research institutions (such as CIRAD, Ifremer, etc.) and higher education institutions (such as universities, *grandes écoles*,…). These institutions take part in social structures called *laboratoires* (referred to as research labs in the following sections). Their financial contributions to the operation of research labs may be multiple, *e.g.* they may pay the salaries or stipends of its members, provide fixed assets such as buildings or infrastructures, pay for resources such as supplies, electricity, etc. A typical research lab benefits from the involvement of several public institutions. It comprises between ten and at most a few hundred members, and occupies one or several buildings. France counts over 1000 public research labs overall. According to the French legislation, all legal public entities comprising at least 250 staff members are bound to build their GHG inventory and define a mitigation action plan every three years, following a pre-defined methodology [MEEM, 2016]. Research labs, be they below or above the 250 member-threshold, do not have to comply with this legislation.

Still, research labs are decision-making entities. For example, experimental designs, scientific goals, as well as access to research facilities are decided and managed at the research lab scale ; a fraction of the annual budget is also managed at lab scale. Consequently, the research lab is a relevant scale to tackle the question of the research’s carbon footprint. *GES 1point5* is specifically designed to address this question and meet the following goals. At the research lab scale, one should be able to analyse the carbon footprint to understand (i) the main emission sources and their relative contributions, (ii) the relative contributions of its members according to various groupings such as seniority or discipline. One should also be able to make decisions to reduce the carbon footprint of the lab and monitor the actions implemented over time. For example, contrasted mitigation decisions or policies may be experimented at lab scale to reduce emissions from professional travel (e.g., an internal carbon tax or individual emission quotas).

At a smaller scale, *GES 1point5* may also allow to estimate the carbon footprint of specific research projects or research teams within the lab. At a larger scale, *GES 1point5* enables an extrapolation to assess the overall carbon footprint of the research sector, and to describe the distribution of emission sources across localities or disciplines.

## 3 *GES 1point5* methodology

To operate on a common ground, *GES 1point5* is designed to comply with the official GHG inventory methodology. It is all the more relevant as the French legislation abides by the GHG Protocol standard [WRI and WBCSD, 2004], which is one of the most used standards in the world. *GES 1point5* is specifically adapted to the context of research labs both in its estimation of carbon footprints and methodological choices (for example it tackles the partitioning between teaching and research). Therefore, *GES 1point5* may be virtually used by research centers in any country. In the next two subsections, we present the scope of *GES 1point5* inventory, *i.e.*, the GHG emission sources considered, as well as the key hypotheses relative to emission factor values.

Methodological choices are comprehensively presented in the *GES 1point5* online documentation.

### 3.1 Scope

The GHGs considered in *GES 1point5* are those of the Kyoto Protocol. *GES 1point5* takes into account the most common and often predominant emission sources in research labs: buildings (through energy consumption for heating, electricity, and refrigeration processes), purchase of digital devices, commuting, and professional travel (due to the use of cars, trains, ships or planes to attend meetings or for field work).

### 3.2 Emission factors

Emission factors represent the amount of GHG emissions (expressed in carbon dioxide equivalent, CO_2_e) generated by a unit of activity. GHG emissions are estimated as follows: for each source of emission, the amount of activity is collected or estimated, and then multiplied by the corresponding emission factor. For example, the emissions of a gasoline-fueled car over a year are calculated as the product of the kilometers traveled over the year by the emission factor of one kilometer traveled.

The emission factors considered in *GES 1point5*, as well as their uncertainties mainly stem from the official ADEME database^13^, which is specifically adapted to the French context. The ADEME database generally includes several types of emission factors for one source of emission. In the case of air travel, four types of emission factors are provided per passenger-kilometer traveled: a factor related to engine fuel combustion, a factor related to upstream emissions from fossil fuel production, a factor related to aircraft manufacture, and a factor related to condensation trails or contrails (*i.e.*, fugitive emissions). *GES 1point5* presents air-travel emissions both with and without condensation trails. Following [Lee et al., 2021], 2/3 of the net warming effects of aviation are due to non-CO_2_ effects, in which contrails contribute most. Various actors do not include the contrails of air flights. This is the case of the International Civil Aviation Organisation^14^. French transportation companies, which are compelled to publish the GHG emissions generated by their services, also generally do not include the contrails of air flights [Ministère de la transition écologique et solidaire, 2018]. By default, *GES 1point5* carbon footprint table posts air travel emissions without contrails in order to comply with the French legislation applied to transportation companies. A note added on top of the table posts the emissions with contrails.

When emission factors are missing in the ADEME database, *GES 1point5* relies on emission factors provided in the most recent available literature together with conservative estimates for associated uncertainties. Furthermore, the interdisciplinary *GES 1point5* team has created customized emission factors to take into account specific research lab activities. For example, the application includes an emission factor for research campaigns at sea.

Regarding digital devices, our emission factors stem from a tool developed by EcoInfo - a group focused on the footprint of digital technologies^15^. The tool is named Ecodiag^16^ and emission factors include emissions from manufacturing (raw material extraction, assembly, packaging) and distribution. They are retrieved from the figures provided by manufacturers, or from default average values when the precise model is not provided or known. Emission factors are estimated for basic configurations, and for lack of a better solution, configuration options such as more powerful CPU/GPU or additional RAM and hard drives are ignored. Emissions from the on-site electricity consumption of devices are already included in buildings emissions. Emissions from digital devices outside of the laboratory (e.g. cloud servers, calculation platforms) are not considered in *GES 1point5* as they are out of the scope of this version.

*GES 1point5* stores around 1,000 emission factors including those of more than 600 district heating systems installed in France. They are updated once a year. For example, those listed as 2020 are obtained from ADEME’s carbon database at the beginning of 2021. The estimation of GHG emissions in *GES 1point5* is based on emission factor values that are the closest in time to the year of the inventory. For emission factors that are bound to change on a yearly basis, typically because the energy mix changes each year (e.g. electricity, district heating), we use the emission factor that relates to the year considered in the inventory (or the closest year if the year’s emission factor is not yet available in the ADEME carbon database). For example, the emission factors for district heating available in the ADEME carbon database in December 2021 are those for 2018. Therefore, for the 2018, 2019 and 2020 GHG inventories, the calculations currently rely on 2018 emission factors.

To adapt *GES 1point5* in a country other than France, a few emission factors need to be adjusted. In particular, emission factors relating to electricity consumption (*i.e.*, the carbon intensity of the national electric grid) and its derivatives like electric vehicles (trains, cars, bicycles) depend on the electricity generation mix of the country. The range of carbon intensities can vary by a factor of 10 or more between countries. The carbon-intensity was 79 gCO_2_/kWh in France in 2013, whereas the United States, China and India reached 522, 766, and 912 gCO_2_/kWh respectively the same year. (Source IEA, 2013. CO_2_ emissions from fuel combustion - hightlights, retrieved from ADEME carbon database in January 2022). To ease the adjustment of the emission factors, they are stored within the same directory in an editable file format.

## 4 Implementation

*GES 1point5* has been implemented as a single-page web application. On the frontend side, the VueJS framework^17^ and the Buefy library^18^ have been chosen to build and design the user interface. On the backend side, the python-based django framework^19^coupled with the django REST library have been used to create the application programming interface (API) that interacts with the database. The application uses input information that can be gathered reasonably easily (provided support is granted by the administrative services) and converts it into a GHG footprint. For each emission source considered in *GES 1point5*, the tool converts GHG-emitting activity levels into CO_2_e, using emission factors as described in Section 3.2.

From its home page (Figure 1), the application offers its users the opportunity to estimate GHG emissions anonymously or using an authenticated account. In the latter case, *GES 1point5* allows to store inputs in a database, described in Figure 2, and provides outputs figures and tables as described in the following sections.

**Figure 1:**
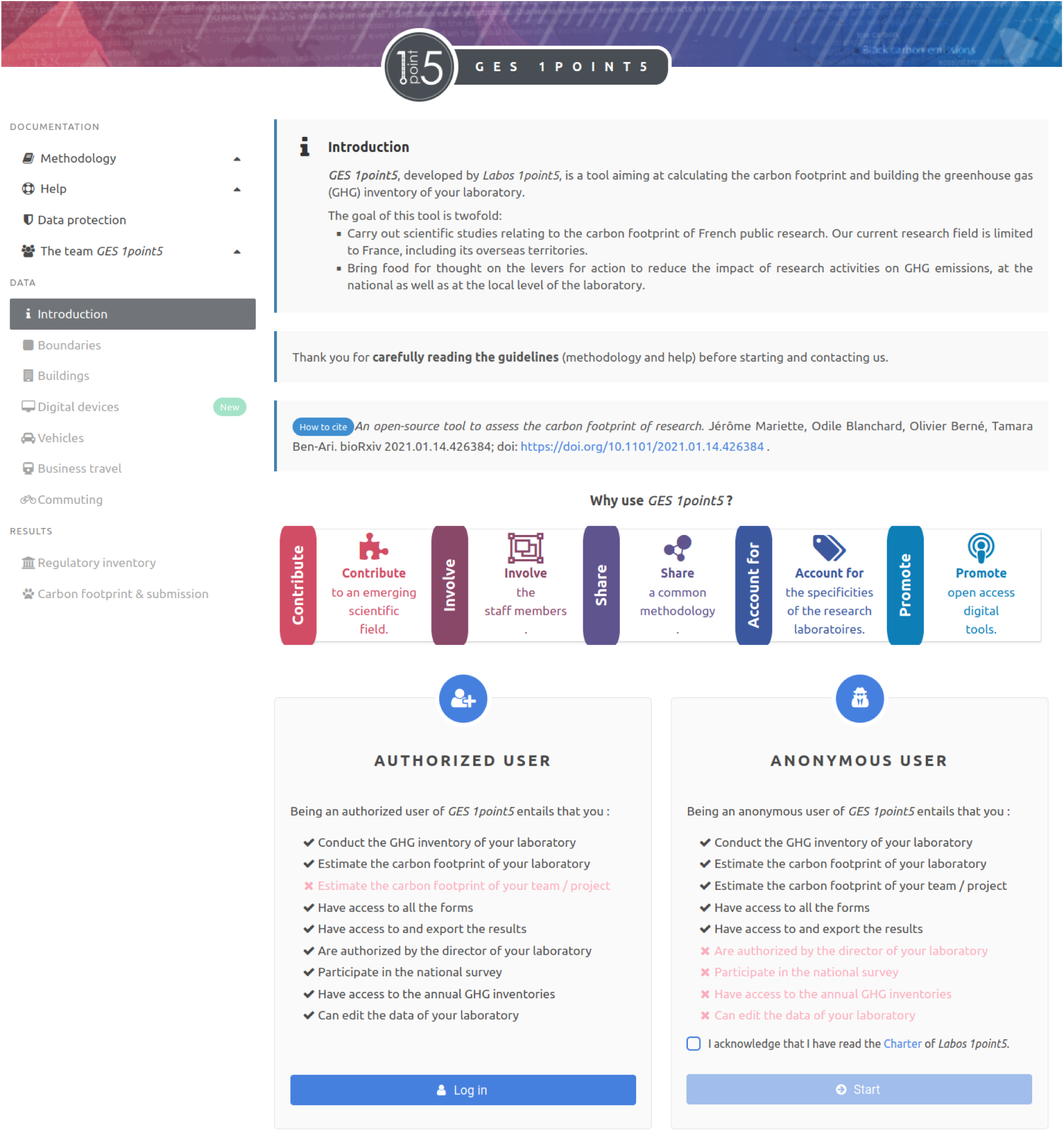
*GES 1point5* home page offers its user the opportunity to estimate GHG emissions anonymously or using an authenticated account. The application menu, on the left, allows to navigate between documentation pages and the different emission sources forms.

**Figure 2:**
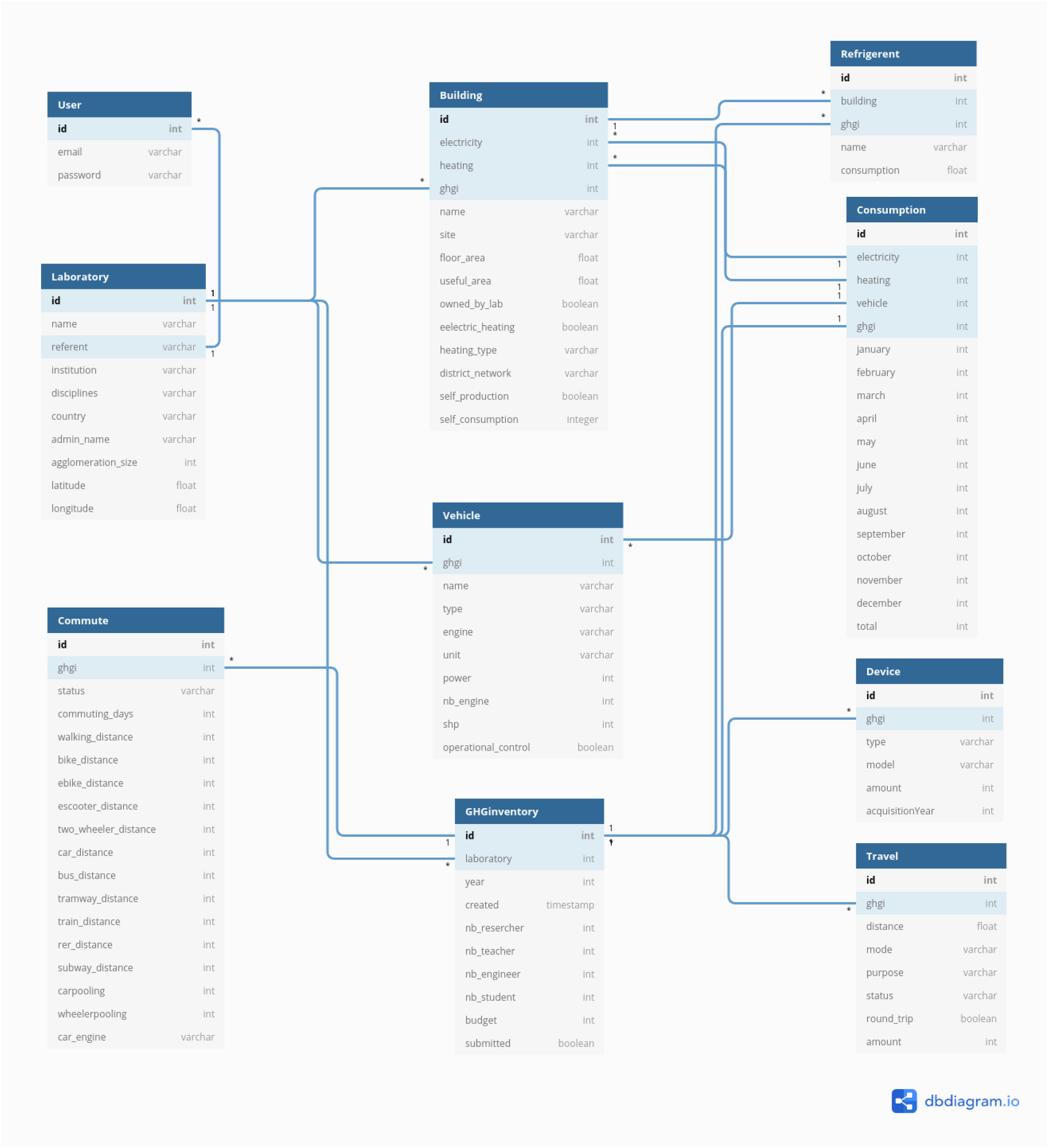
*GES 1point5* database diagram allowing to store input data collected by forms and routines.

### 4.1 Inputs

To gather the data required as inputs, *GES 1point5* provides a set of forms and routines briefly explained below.

- General information: year of the GHG inventory; number of lab members according to their position (i.e. researchers, professors, engineers, PhD students and post-doctoral fellows). These data are useful to perform statistical analyses of the emissions across the various positions, e.g. emissions from commuting, or from traveling.
- Buildings: floor area ; electricity consumption, heat, and refrigerant gases; particularities related to the generation of electricity, when applicable (e.g. use of solar panels). The floor area allows to compute emission ratios per square meter. When a laboratory covers part of its electricity consumption through self-generation, emissions relate to net consumption, *i.e.* the consumption metered through the power company minus self-consumption.
- Vehicles operated by the laboratory: type (e.g. car, motorcycle, aircraft); type of fuel; power, distances traveled, number of hours of operation when applicable. These data allow to compute the energy consumed by the various vehicles. Figure 3 presents the form dedicated to adding a new vehicle and entering the features needed to calculate its energy consumption.
- Digital devices purchased by the laboratory : type (e.g. desktop, server, monitor, video projector, smartphone, printer, wifi hub); brand name ; model. These data allow to allocate to each device its specific emission factor as described in Section 3.2.
- Commuting: a standardized online survey dedicated to collecting the commutes of the lab members is embedded in *GES 1point5*. The survey is sent to all lab members and collects the number of commuting days per week, the modes of transportation used and the distances traveled for one or two frequent standard commutes. The data are automatically imported in *GES 1point5* and corrected for non-response by the number of lab members according to their academic position. To raise the respondent’s awareness, *GES 1point5* offers an estimate of the respondent’s annual commuting emissions at the end of the survey, as presented in Figure 4. From this display, the respondent can simulate different commuting scenarii and thus estimate emission reduction options. Once the survey period is over, *GES 1point5* provides a curation tool to analyze the survey results and remove potential extreme answers as shown in Figure 5.
- Professional travel: raw data are extracted from the information systems of the various research institutions that pay for the trips of lab members. For each trip, they include information on date, departure and destination places (cities,countries), travel modes, travel purpose and lab member position. These data are imported as a tab-separated values (.tsv) file into *GES 1point5*. Table 1 defines the required format. For each leg of a trip, collected data allow to calculate traveled distances and multiply it by the emission factor of the transportation mode used.

**Figure 3:**
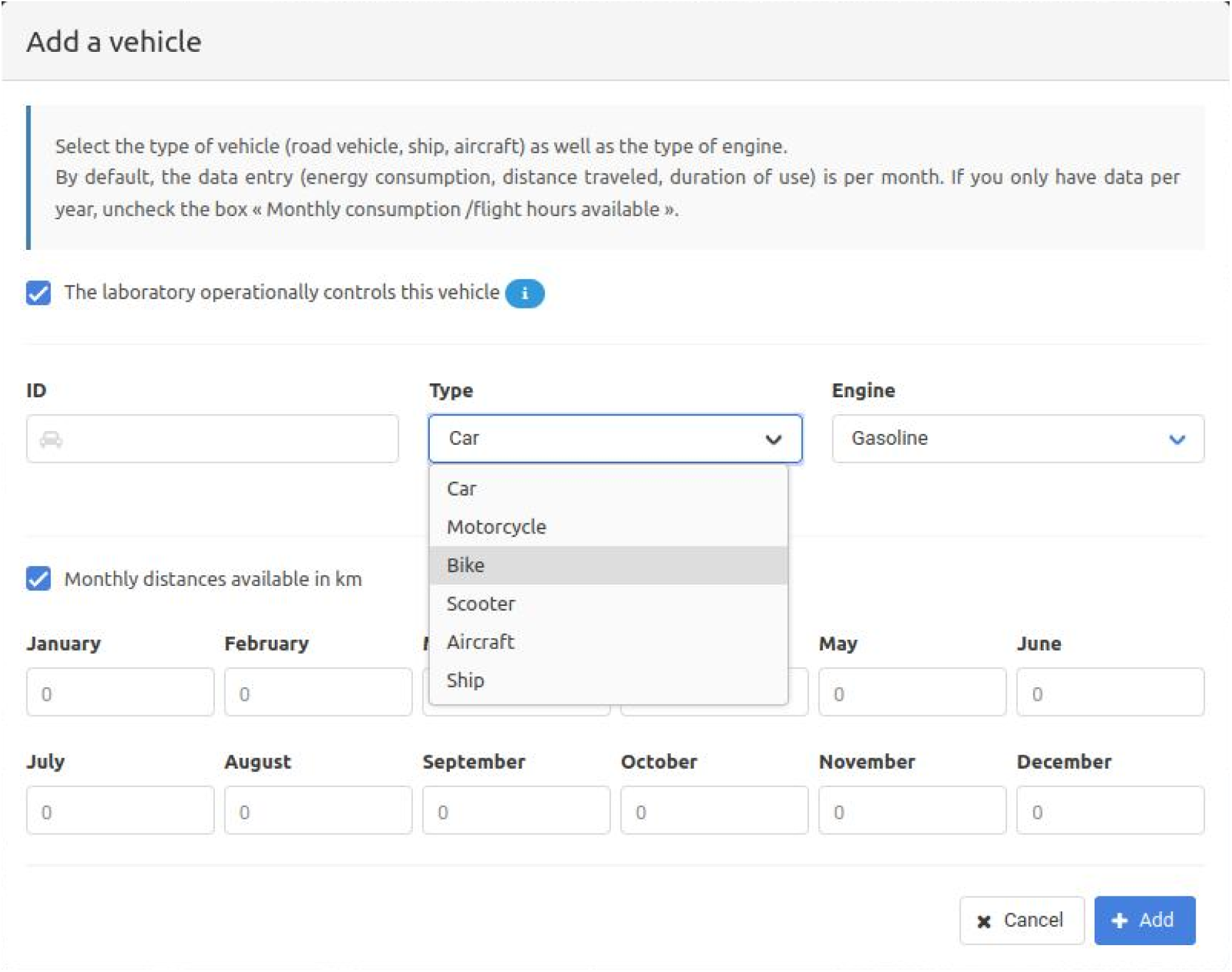
Form to add a new vehicle to the inventory of the research lab. The vehicle can be either a car, a motorcycle, a bike, a scooter, an aircraft or a ship. The form requires to define the vehicle motorisation, its annual energy consumption or the number of hours / days of operation.

**Figure 4:**
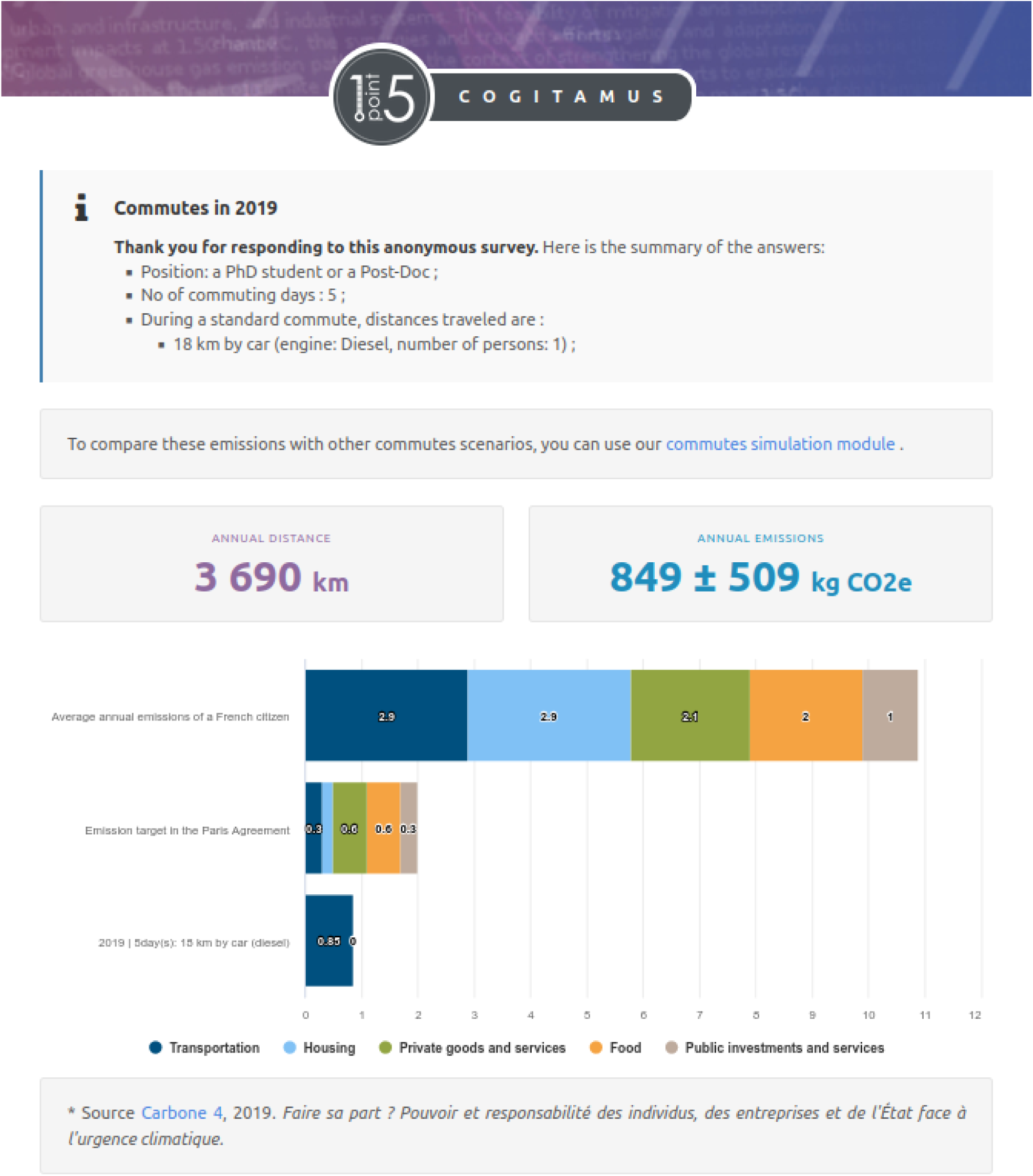
To raise lab members’ awareness, *GES 1point5* provides an estimate of the respondent’s annual distance traveled and commuting emissions at the end of the survey. These emissions are compared to the average annual emissions of a French citizen and to the emission target needed to comply with the Paris Agreement.

**Figure 5:**
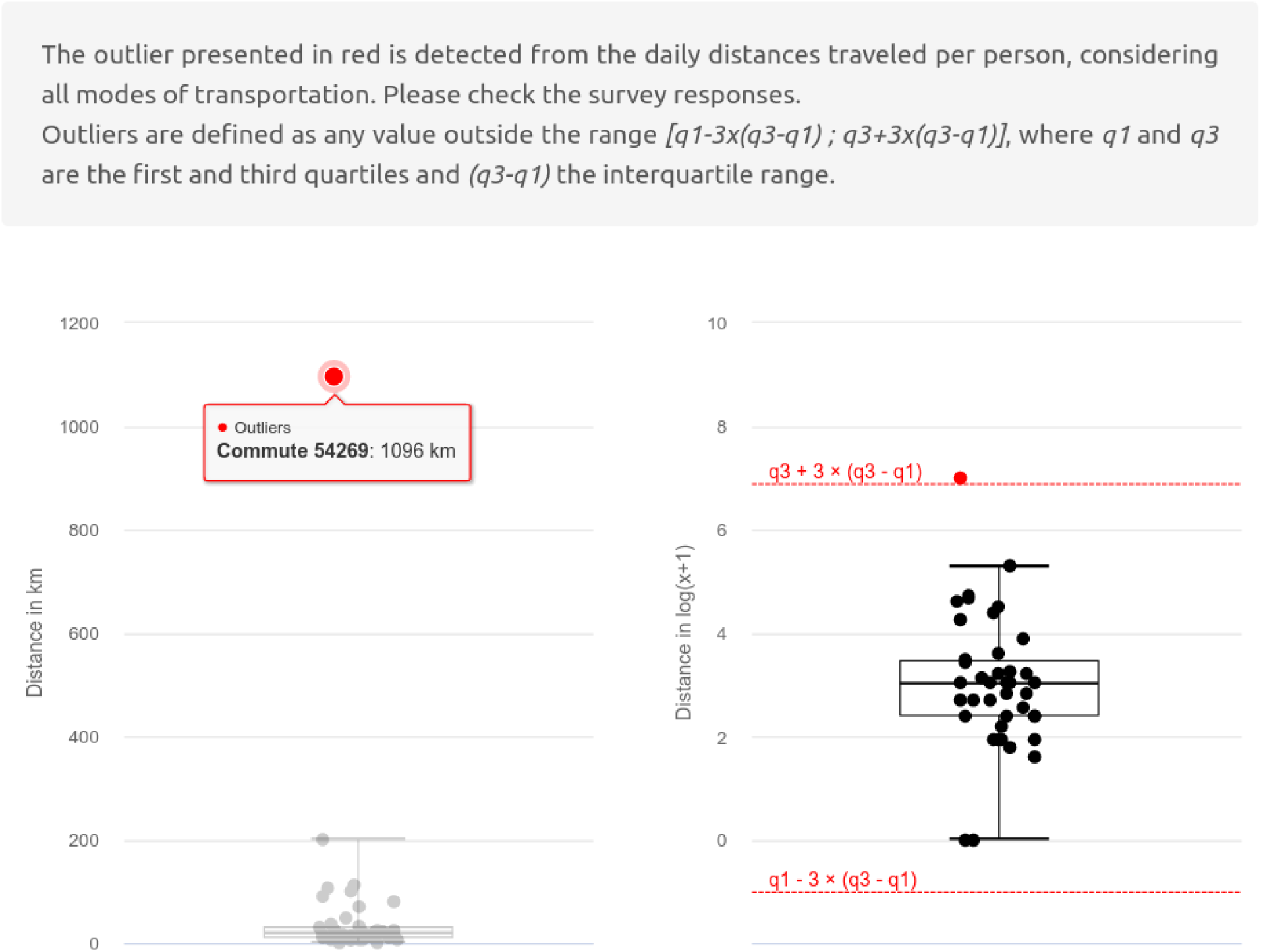
*GES 1point5* curation tool to analyze the survey results and remove potential extreme answers.

**Table 1:** Column description of the .tsv file accepted by *GES 1point5* to import professional trips relating to business travel.

| Column ID | Description |
| --- | --- |
| Trip number | A trip is a set of legs. The trip number is a simple sequence number (from 1 to n) which allows to gather all the legs of the same trip. One line per leg of the same trip. If the return trip is different from the outward trip, extra lines must be created. |
| Departure date | A date in dd/mm/yyyy format. |
| Departure city | The departure city name. This field is used to obtain the departure city geographic coordinates using the *geonames*<sup>21</sup> database. |
| Departure country | The departure country or its ISO3166 code. |
| Destination city | The destination city name. This field is used to obtain destination city geographic coordinates using the *geonames* database. |
| Destination country | The destination country or its ISO3166 code. |
| Travel mode | This field can take a value among [ "Plane", "Train", "Car", "Taxi", "Bus", "Tramway", "Paris suburban express railway including RER", "Subway", "Ferry" ]. |
| Number of people in the car | In case of a trip in a car or a taxi, number of people in the vehicle. |
| Roundtrip | If the outward and return trips are identical, enter "YES", otherwise "NO". |
| Travel purpose (optional) | This optional field allows the application to perform emission statistics based on the travel purpose. This field can take a value among [ "Field study", "Conference", "Seminar", "Teaching", "Collaboration", "Visit", "Research management ", "Other" ]. |
| Agent position (optional) | This optional field allows the application to perform emission statistics based on the agents' position. This field can take a value among [ "Researcher or professor", "Engineer, technician or assistant", "PhD student or Post-Doc", "Guest" ]. |

No personal data as defined by the European General Data Protection Regulation^20^ (GDPR) is stored by GES 1point5. The commuting survey complies with the GDPR. Professional travel data are collected anonymously. Furthermore *GES 1point5* abides by strict data minimisation rules : it does not store the origin and destination locations of the professional trips; it only stores travel distances.

### 4.2 Outputs

As mentioned above, *GES 1point5* complies with the French legislation, which itself abides by the GHG Protocol standard [WRI and WBCSD, 2004]. *GES 1point5* thus provides scope 1 (direct emissions from owned or controlled sources) and scope 2 (indirect emissions from the generation of purchased electricity, heating and cooling) results, as well as the following emission categories among all other indirect emissions of scope 3: energy-related emissions not included in scope 1 or 2, fixed assets, business travel, employee commuting. The resulting table, presented in Figure 6, can be downloaded.

**Figure 6:**
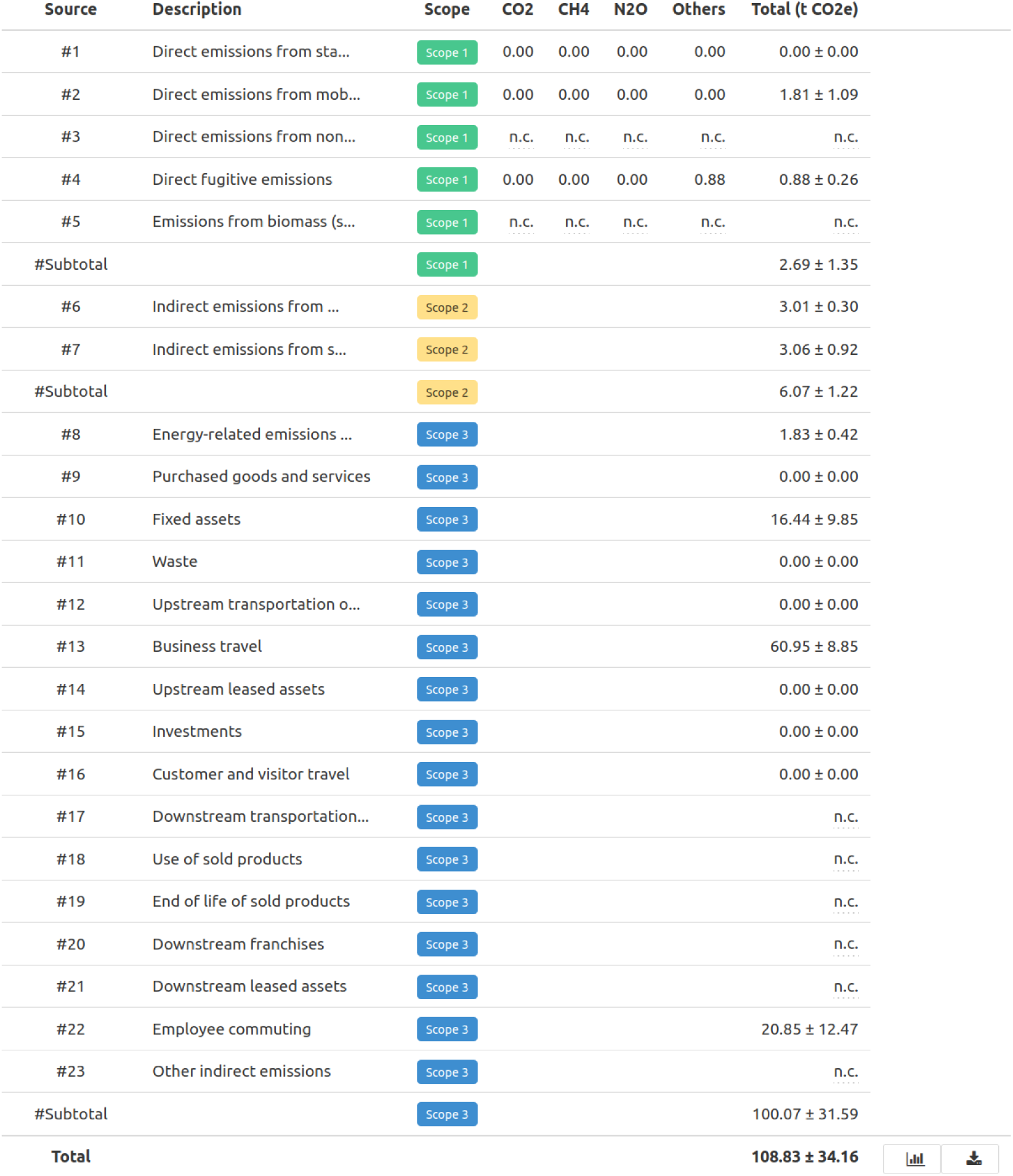
Illustration of a regulatory table obtained with *GES 1point5* for a fictitious research lab, presenting the emissions distributed among the three scopes of the GHG Protocol standard. *n.c.* is displayed for emission sources that are not computed by *GES 1point5*.

In addition to displaying GHG emissions distribution within the three regulatory and GHG Protocol scopes, *GES 1point5* provides a user-friendly synthetic table and a graphical representation of carbon footprints (*i.e.*, GHG emissions are aggregated per activity). This additional representation is designed to help users to identify predominant emission sources and decide which actions to implement in order to efficiently mitigate emissions. For example, while direct and indirect emissions generated from buildings’ heating and cooling systems are split among the three scopes in the regulatory display, they are aggregated in the carbon footprint representation. Similarly, the travel carbon footprint aggregates GHG emissions of vehicles, commutes and professional travels. A few additional examples are provided in Figures 7 and 8. Finally, when users have entered all input data and therefore obtained regulatory and carbon footprint estimates and corresponding results in a downloadable format, they are invited to submit their data. Data are then stored in *GES 1point5* database, as described in Figure 2. The data submitted contribute to the building of a national database aggregating hundreds (as of 2022) of carbon footprints from a large variety of research labs in France, allowing in depth research on the carbon footprint of the French public research sector as explained in Section 2.

**Figure 7:**
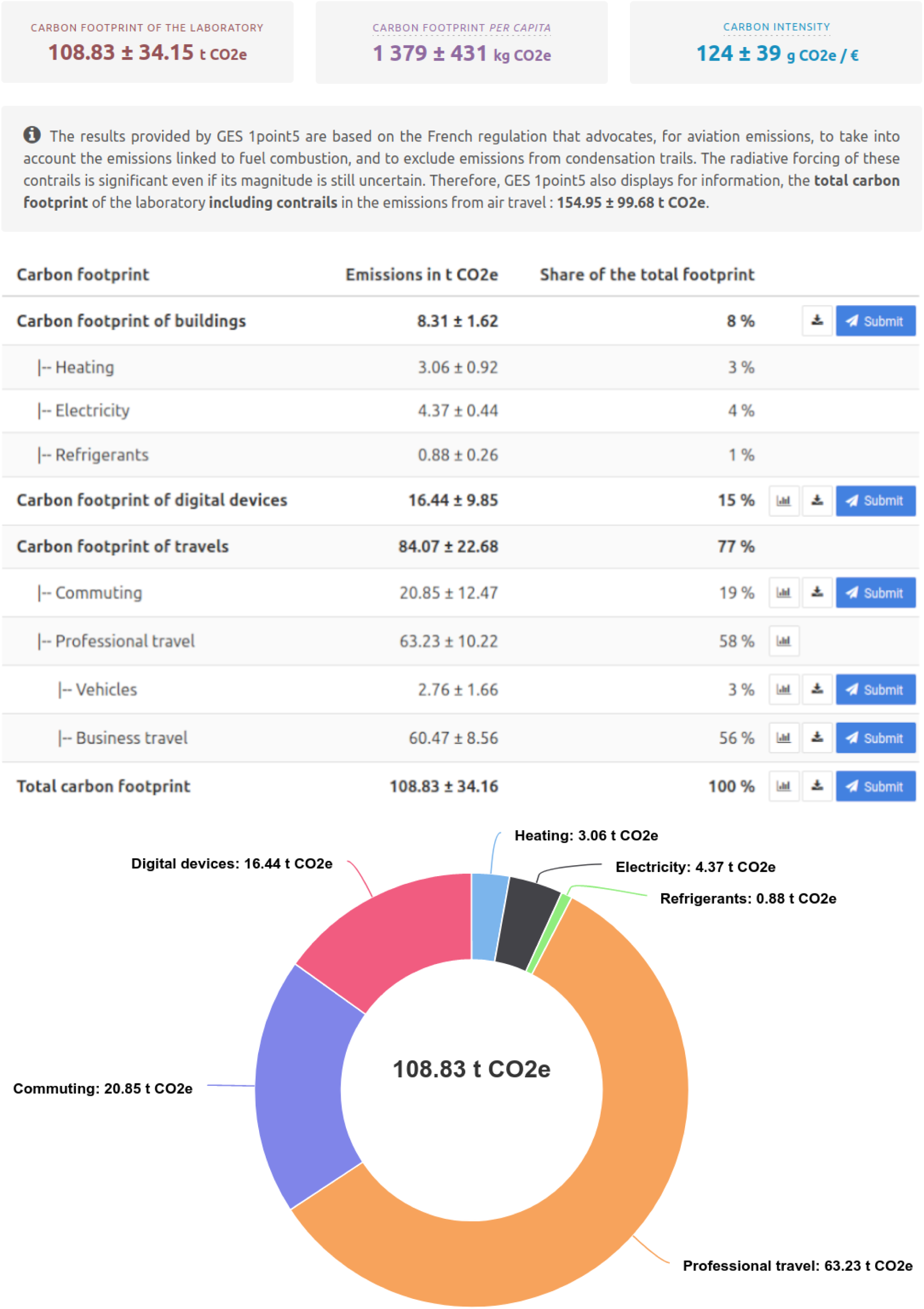
Illustration of the carbon footprint information provided as an output of *GES 1point5* : table (upper panel) and pie chart (lower panel), showing the distribution of emissions in tons of CO_2_e.

**Figure 8:**
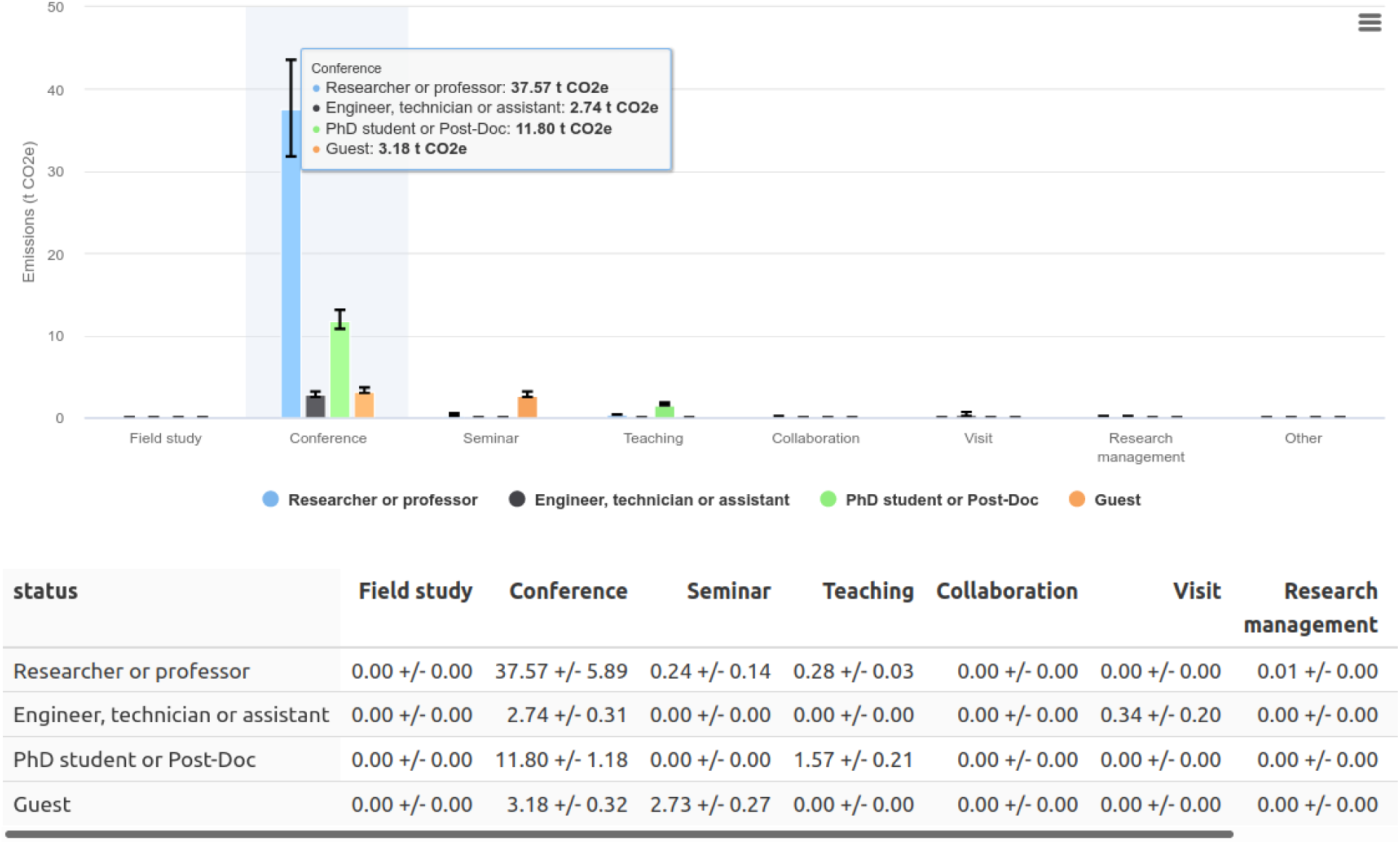
Illustrative distribution of the professional travel carbon footprint provided by *GES 1point5* according to travel purposes and professional position.

## 5 Results

### 5.1 Simulated data and results for a research lab

This section provides and discusses results of the 2019 carbon footprint of a fictitious research lab named Cogitamus. This example does not aim at providing a representative case study since there is a large heterogeneity between research labs, but at illustrating *GES 1point5*, including its visual outputs. Cogitamus fictitious lab comprises 80 members distributed as follows: 14 researchers, 24 associate professors, 17 engineers or administrative staff and 25 PhD students or postdoctoral fellows. Its 2019 budget amounts to 875 500 Euros. Cogitamus occupies one building shared with another lab and 60% of the total floor space, 3,300 *m*^2^. In 2019, the fictitious building consumed 200,000 kWh LHV (lower heating value) from the Toulouse Canceropole urban heating network, 120,000 kWh of electricity and 0.3 kg of the R32 refrigerant gas. Cogitamus also owns one diesel car, which traveled 12,000 km in 2019. For a full reproducibility of the results presented in this section, the fictitious commuting survey results^22^, professional travel file^23^ and digital devices inventory^24^ are freely available.

Figures 6, 7 and 8 illustrate the GHG emissions inventory for Cogitamus that complies with the French regulation and the GHG Protocol standard, the user-friendly carbon footprint representation, graphs excerpted from *GES 1point5*, respectively. More specifically, figures 6 and 7 display Cogitamus GHG emissions (108.83 *±* 34.16 t CO_2_e) from two different perspectives, *i.e.*, the regulatory GHG inventory table and the detailed carbon footprint. The latter representation also provides the lab total carbon footprint including the effects of contrails of air travel (154.95 *±* 99.68 t CO_2_e), the *per capita* carbon footprint (1 379 *±* 431 kg CO_2_e) and the carbon intensity of Cogitamus (124 *±* 39 g CO_2_e / €). Figure 7 shows that Cogitamus GHG emissions are mainly driven by professional travel (58%), followed by commuting (19%) and digital devices (15%). Within professional travel, trips to conferences prevail Figure 8.

### 5.2 Database for a network of labs

Since the public release of *GES 1point5* in October 2020, more than 610 GHG inventories have been initiated by 395 different research labs. This dynamic, represented in Figure 9, shows the growing interest of labs in monitoring their carbon footprint as well as the software learnability and operability by non-specialists. The data thus collected contribute to the creation of a unique database permitting to identify robust determinants of research carbon footprint.

**Figure 9:**
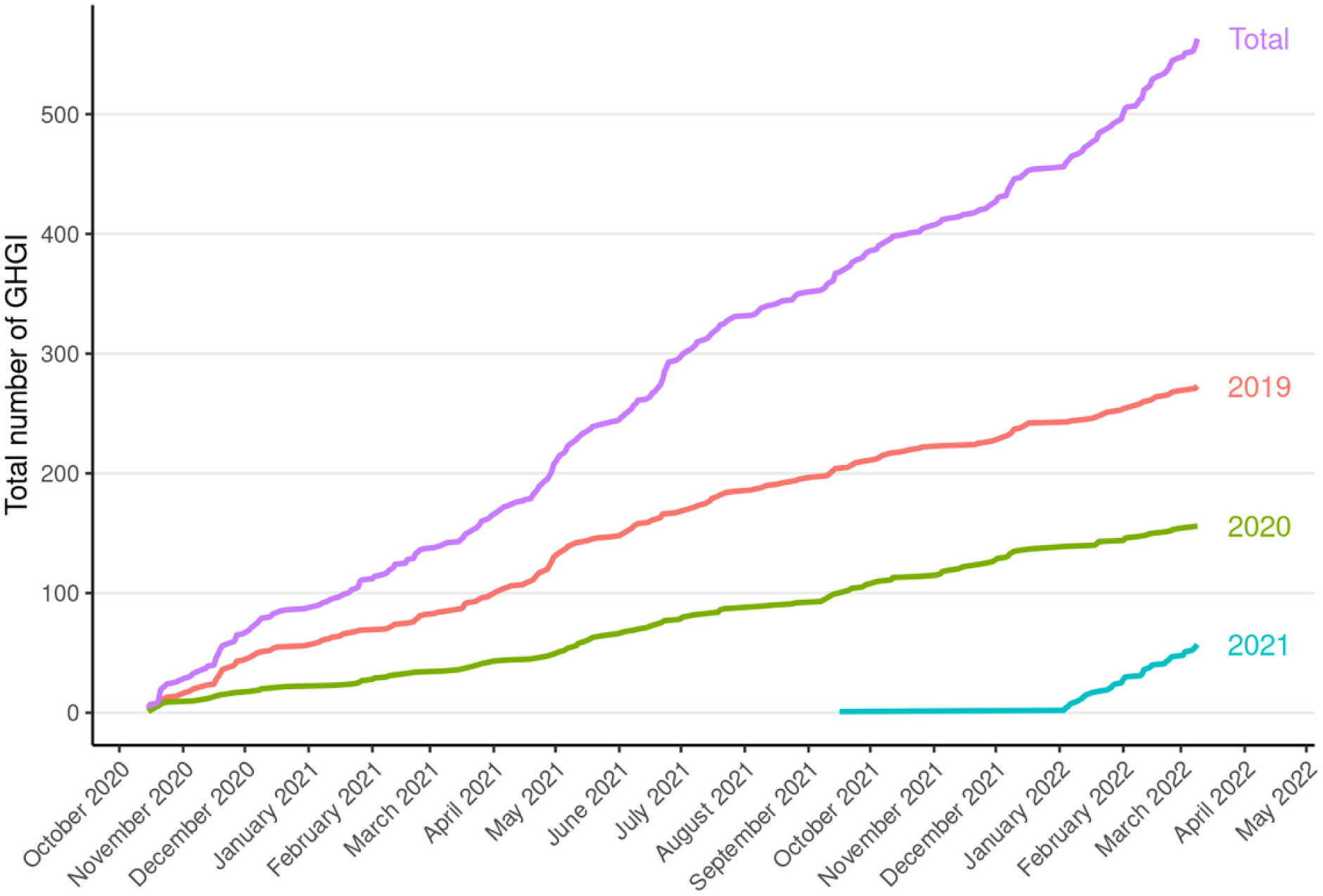
Evolution of the creation of annual GHG inventories in *GES 1point5* for the years 2019, 2020 and 2021.

Thus far, 108 GHG inventories have been submitted for 2019 into *GES 1point5* database, which means that they are complete and preliminary analysis can be carried out. Considering the current scope of *GES 1point5*, the average emissions are 591 t CO_2_e for a research lab and 4.5 t CO_2_e for a lab member. Figure 10 shows the share of each source. The great heterogeneity observed between research labs suggests that there will not be a *one-size-fits-all* emission mitigation solution for all the laboratories, but customized solutions tailored to the specificities of the various laboratories.

**Figure 10:**
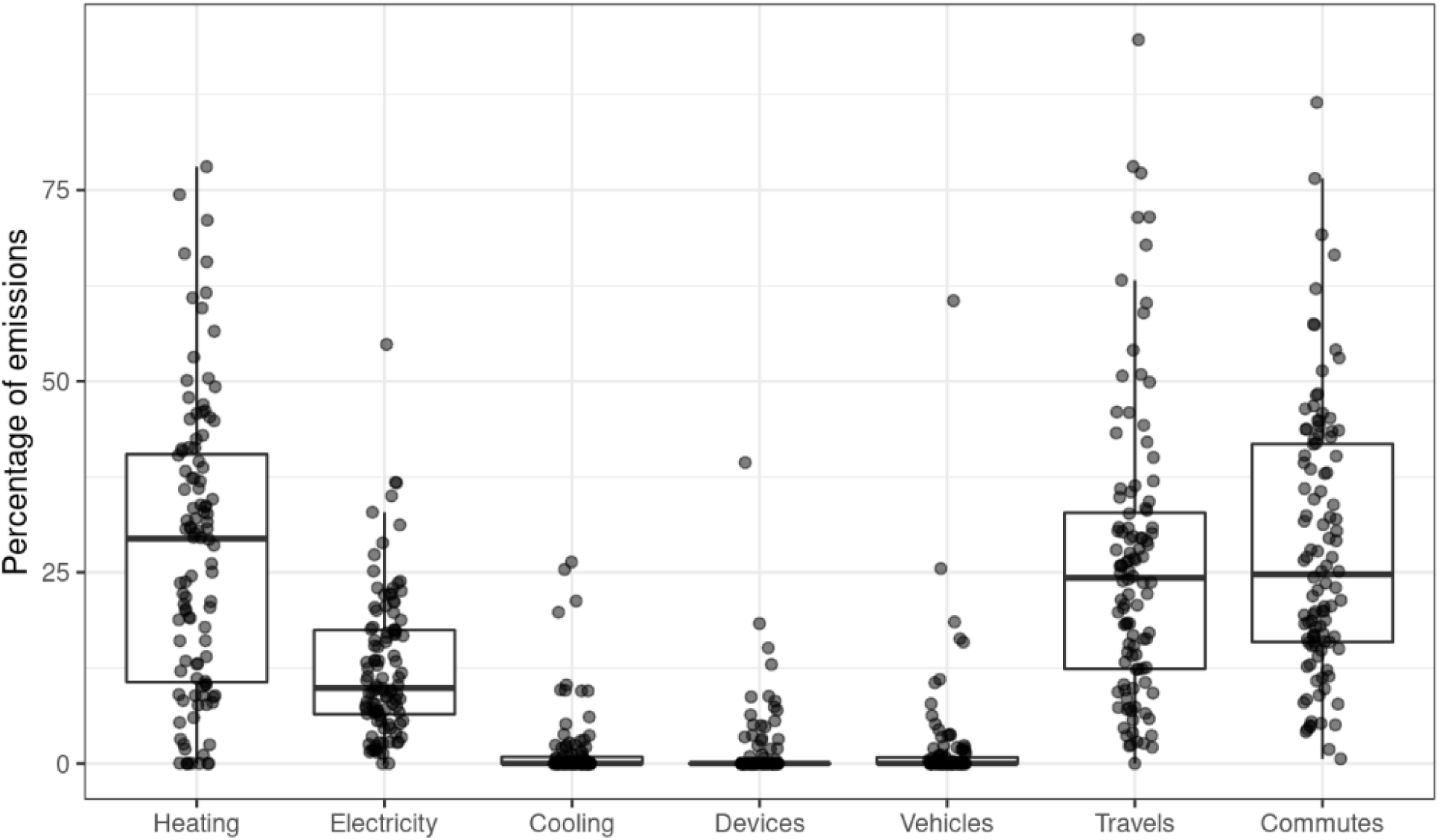
Percentage of the GHG emissions by sources for the 108 GHG inventories submitted in *GES 1point5* for the year 2019. For each emission source, each dot represents a research lab.

## 6 Discussion and conclusion

*GES 1point5* is a first step in a larger endeavour to inform on, facilitate, and enable emission reductions from research activities in France and at the international level. While an alternative lies in the use of external consultants both in the estimation of GHG emissions and on the design of reduction options, *GES 1point5* is developed to help the research community to tackle the problem by itself. It is designed to increase expertise and to provide the community with a free and transparent tool to facilitate broad adoption and mitigation actions. *GES 1point5* is one of the building blocks of the *Labos 1point5* project. *Labos 1point5* is currently working on the design of a network of research labs in transition to low-carbon practices. *GES 1point5* will have a pivotal role in the design and monitoring of these changes.

The current version of *GES 1point5* focuses on common emission sources that are often predominant in research labs. The next version of *GES 1point5* will address emissions linked to purchased goods other than those already taken into account in the tool. Note that the estimation of the carbon footprint of purchased goods is more complex because most needed emission factors do not exist. For example, it is frequent to use specific chemical solvents in biology labs, with only very little information on the manufacturing and supply chains. Various possible methodologies are currently under study and *Labos 1point5* thrives to consider the right trade-off between comprehensiveness, representativeness and data accessibility. *GES 1point5* will also aim at estimating the GHG footprint of large scale scientific infrastructures (e.g. clusters and big supercomputers, particle accelerators, telescopes, etc.) that provide research services to multiple research labs. The goal is to estimate emission factors per unit of service provided so that the emissions generated by these large infrastructures may be distributed to the labs using the services. Finally, future versions of *GES 1point5* will aim at providing a scenario-building tool to explore possible futures in order to inform decisions and decide on actions to be implemented. This work is part of the *Labos 1point5* research project.

The research sector must reduce the carbon footprint of its activities along with all other sectors, in order for France to reach the goal of the Paris Agreement. One may also argue that the responsibility of academia is to lead this transformation by “walking the talk”. A first step naturally pertains to estimating the current level of emissions. In this context, a standardized tool is essential: it paves the way for a deeper understanding of the main drivers of emissions and of the influence of discipline or local specificities; it enables the tailoring of coordinated mitigation strategies, as well as the reporting and monitoring of those actions. *GES 1point5* is designed to address these needs.

As an open-source software, *GES 1point5* is freely accessible and replicable around the world, provided emission factors are adjusted to the country. Users from research institutions around the world may find a real value added in *GES 1point5*, as it is an online tool that presents outputs not only in the GHG Protocol inventory format, but also under the more operational format of carbon footprints. This will enable a fruitful comparison between research centers worldwide to get a deeper understanding on the barriers and levers to change.

## 7 Acknowledgment

We acknowledge Géraldine Sarret, Maximilien Chaumon, Emmanuel Faure, Iago Bonnici, Arthur Leblois, Maxence Morel, Xavier Schwindenhammer and Marie Stephane Trotard for their help and feedback. We thank L’association le PIC^25^ for support on technical aspects related to *GES 1point5*. The work presented in this article was conducted as part of the *GDR Labos 1point5* with financial support from ADEME, CNRS and INRAE.

## Footnotes

1 https://www.unh.edu/sustainability/research/campus-calculator-tools

2 https://ghgprotocol.org/

3 https://www.associationbilancarbone.fr/les-solutions/

4 https://co2.myclimate.org/en/

5 https://www.carbonfootprint.com/

6 https://offset.climateneutralnow.org/footprintcalc

7 https://nosgestesclimat.fr/simulateur/bilan

8 https://www.goodplanet.org/fr/calculateurs-carbone/particulier/

9 http://labos1point5.org/ges-1point5

10 *GES* stands for *GHG* in French

11 https://labos1point5.org/

12 https://www.sciencemag.org/careers/2006/04/finding-your-way-around-french-research-system

13 https://www.bilans-ges.ademe.fr/en/accueil/

14 https://www.icao.int/environmental-protection/Carbonoffset/Pages/default.aspx

15 https://ecoinfo.cnrs.fr/

16 https://ecoinfo.cnrs.fr/ecodiag-calcul/

17 https://vuejs.org/

18 https://buefy.org/

19 https://www.djangoproject.com/

20 https://eur-lex.europa.eu/eli/reg/2016/679/oj

22 https://cloud.le-pic.org/s/kWi6PLPRo9PRZFo

23 https://cloud.le-pic.org/s/TGgCAD2jeyK8JQB

24 https://cloud.le-pic.org/s/enqXfNABAMFJRNs

25 https://www.le-pic.org/

## Notes

https://framagit.org/labos1point5/l1p5-vuejs

### Competing Interest Statement

The authors have declared no competing interest.

